# The effect of macromolecular crowding on single-round transcription by E. coli RNA polymerase

**DOI:** 10.1101/218321

**Authors:** SangYoon Chung, Eitan Lerner, Yan Jin, Soohong Kim, Yazan Alhadid, Logan Wilson Grimaud, Irina X. Zhang, Charles M. Knobler, William M. Gelbart, Shimon Weiss

## Abstract

Biological reactions in the cellular environment differ physicochemically from those performed in dilute buffer solutions due to, in part, slower diffusion of various components in the cellular milieu, increase in their chemical activities, and modulation of their binding affinities and conformational stabilities. *In vivo* transcription is therefore expected to be strongly influenced by the ‘crowdedness’ of the cell. Previous studies of transcription under macromolecular crowding conditions have focused mainly on multiple cycles of RNAP-Promoter associations, assuming that the association is the rate-determining step of the entire transcription process. However, recent reports demonstrated that late initiation and promoter escape could be the rate-determining steps for some promoter DNA sequences. The investigation of crowding effects on these steps under *single-round* conditions is therefore crucial for better understanding of transcription initiation *in vivo*. Here, we have implemented an *in vitro* transcription quenched-kinetics single-molecule assay to investigate the dependence of transcription reaction rates on the sizes and concentrations of crowders. Our results demonstrate an expected slowdown of transcription kinetics due to increased viscosity, and an unexpected enhancement in transcription kinetics by large crowding agents (at a given viscosity). More importantly, the enhancement’s dependence on crowder size significantly deviates from hard-sphere model (scaled-particle theory) predictions, commonly used for description of crowding effects. Our findings shed new light on how enzymatic reactions are affected by crowding conditions in the cellular milieu.

## Introduction

The cellular environment significantly differs from that of dilute buffer. Cells contain many macromolecules in a small volume, resulting in a highly condensed environment(1–3). For example, in *Escherichia coli* (*E. coli*), macromolecules take up to ~40% of the overall volume of the cytoplasm(3). Such a dense environment could alter biological reactions significantly as compared to the same reactions in buffer(4–7). The high viscosity slows macromolecular motions, resulting in slower kinetics(8, 9). In addition, as macromolecules occupy a large fraction of the cellular space, the available volume for other molecules decreases because each macromolecule excludes other molecules from its vicinity. The corresponding excluded volume(6, 10, 11) of a macromolecule is a function of its shape and size as well as those of other nearby molecules(10). This volume exclusion significantly affects the thermodynamics and/or kinetics of reacting molecules, especially when the reaction causes a change in the volume of reactants. It has previously been shown that excluded volume of macromolecules in the crowded environment considerably enhances the kinetics and thermodynamics of the association reactions, and also affects the size of intrinsic disordered proteins, stability of native protein structures, and the folding behavior of proteins and RNAs(12–14).

Transcription is a highly regulated and crucial first step in gene expression, and therefore has been extensively studied(15–19). The great complexity of *in vivo* transcription directed many researchers to take a reductionist approach, focusing on *in vitro*-reconstituted transcription systems in buffer(16, 18, 19). However, this approach sometimes gives rise to results that differ from *in vivo* assays, both quantitatively and qualitatively. These differences are possibly due to environmental factors, such as the crowding effect(20, 21).

Recently, several works reported that transcription reactions could be affected by macromolecular crowding(20, 22–24). For example, studies utilizing cell-free protein expression systems have shown that transcription by T7 RNA polymerase (T7 RNAP) was markedly enhanced under crowding conditions(20, 22). This enhancement was thought to be mainly attributable to increase in affinity between promoter DNA and T7 RNAP as a result of crowding with the assumption that the association is the rate - determining step of the entire transcription reaction(20, 22). However, the effects of crowding on late initiation and promoter clearance are still poorly understood; most studies on transcription under crowding conditions have focused on multiple cycles of transcription, specifically on binding events of DNA and RNAP to form the RNAP-Promoter open complex. Moreover, some recent studies reported that these steps could be rate-limiting for some promoters (e.g. with unique recognition and initial transcribing regions(25, 26)). The detailed understanding of how crowding affects these initiation steps is therefore needed.

In this work, we utilized an *in vitro* single-round quenched kinetics transcription assay using single-molecule detection(26, 27). Our assay is based on the hybridization of a doubly-end-labeled probe DNA oligo to the RNA produced by the transcription reaction. Using this assay, we were able to monitor transcription reactions in various controlled environments that mimic the crowded cellular environment (Figure 1). We studied the effect of the size and concentration of various macromolecular crowding agents on stable open complexes (RP_ICT=2,_ see details in SI), and validated these studies using an RNA-binding-dye assay. We demonstrate that single-round transcription kinetics - starting from the point where RNAP binds the promoter DNA specifically to form an open transcription bubble - are significantly affected by crowders. However, the dependence of the kinetics on crowder size is not consistent with the prediction from the hard-sphere-model based scaled particle theory (SPT), the prevalent framework for elucidating the effect of macromolecular crowding(10, 14, 28).

**Figure 1.**
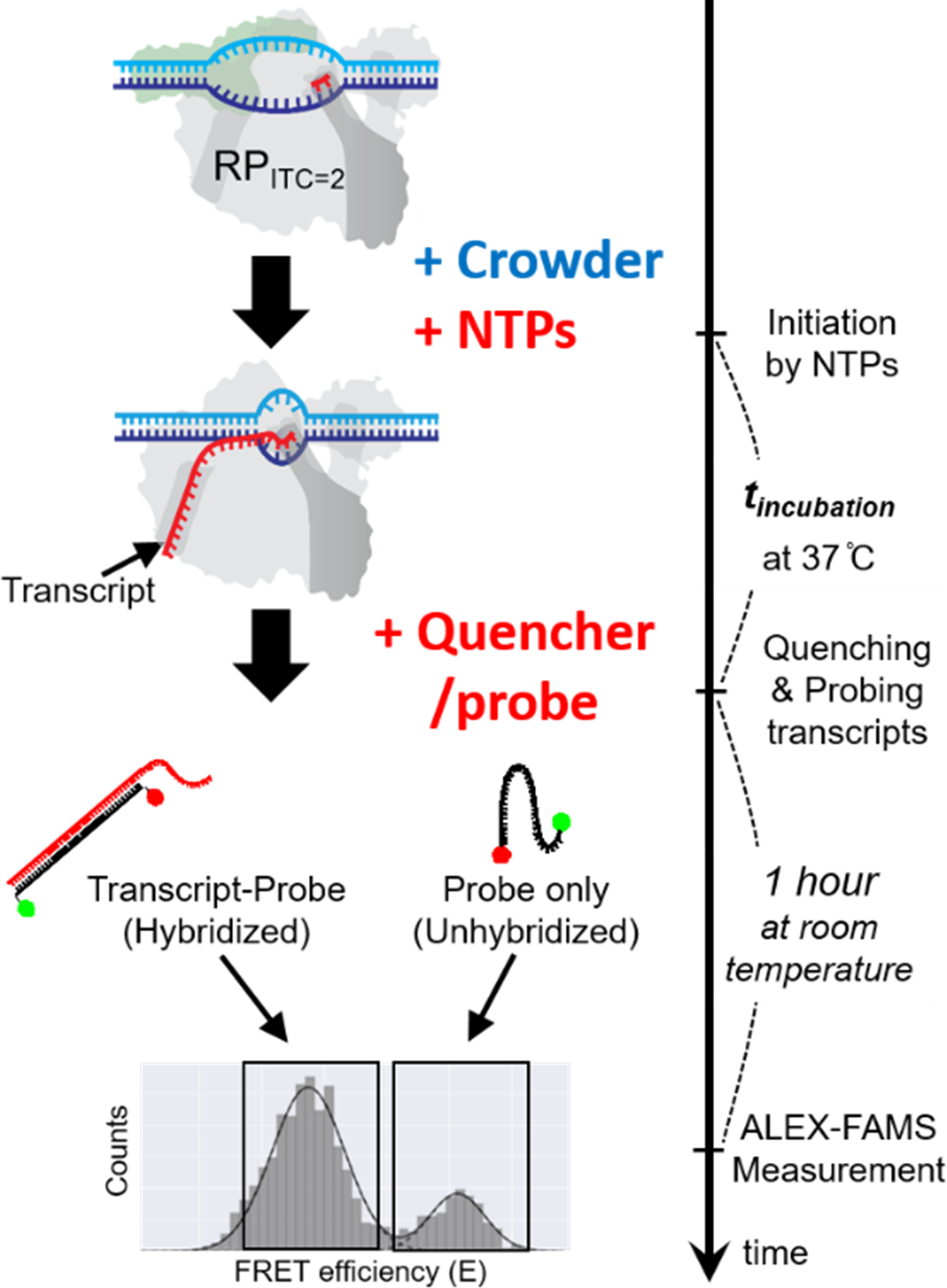
A schematic of the *in vitro* single-round quenched-kinetics transcription assay. The transcription from stable open complexes (RP_ITC=2,_ prepared in buffer) starts with addition of NTPs under crowding conditions and proceeds until quenched by a reaction quencher. RNA transcripts (red) produced by single-round transcription reactions during *t*_*incubation*_ are hybridized with the ssDNA probe (black line with green/red dots) that has sequence complementary to part of the transcript (Fig. S1). The hybridized fraction can be detected by ALEX-FAMS because the stretched structure of hybridized probe shows lower FRET efficiency compared to that of an unhybridized probe(26, 27, 29).

## Results

Our *in vitro* transcription quenched-kinetics assay is a hybridization-based assay that quantifies the amount of transcripts at each time point, defined by the time a reaction quencher is added(26, 27). Once the transcription reaction is stopped by the quencher, the added ssDNA FRET probes hybridize to the transcribed RNAs (during an incubation period). Since the distance between the donor (D) and acceptor (A) dyes on the ssDNA probe increases upon hybridization, a new sub-population of reduced FRET efficiency (E) appears in an ALEX-FAMS 2D histogram – see Figure 1, allowing quantification of the number of transcripts(26, 27).

The transcription efficiency during a given incubation period could be extracted by normalizing the number of single-molecule events for the hybridized sub-population (low E) to the sum of the non-hybridized probe sub-population (high E) and hybridized sub-population (low E). Since the concentrations of probe and RP_ITC2_ are identical in all measurements, a greater fraction of hybridized probe indicates a higher number of transcripts, that is, higher efficiency of transcription for a given time point(26, 27) (Fig. 1).

Single-round transcription kinetics (see Materials and methods #3 for details) starting from stable open complex (RP_ITC=2_), in the presence of various crowding conditions, were measured using the *in vitro* transcription quenched kinetics assay(26) to test whether crowders act through modulation of transcription kinetics or through modulation of the activity of RNAP in transcription (*i.e.* modulating the number of active RNAP-DNA complexes) after RP_ITC=2_ formation. Transcription kinetics were tested with 25% of Glycerol, 15% of PEG 8000, 15% of Ficoll70, and 5% Dextran500 (w/v). Transcription kinetics in buffer (*i.e.* in the absence of viscogen or osmolytes) were also measured as a reference.

Figure 2B shows that the same steady-state levels (t=∞) were reached for reactions in buffer as in the presence of tested osmolytes, regardless of their size and type. This shows that osmolytes do not affect the activity of RNAP. Surprisingly, however, transcription *kinetics* in polymer solutions (containing PEG, Dextran, and Ficoll) are faster than the kinetics in 25% glycerol, even though the viscosities of the polymer solutions are much higher than that of 25% glycerol. Many studies (29–31) have demonstrated that the actual viscosity in a crowded medium (referred to as microviscosity) could differ from the bulk viscosity of the medium. We therefore performed FCS experiments to estimate the actual viscosities (microviscosities) experienced by RP_ITC=2_ for the various crowding environments. These FCS measurements demonstrated that microviscosities for the large crowders Ficoll70 and Dextran 500 are much smaller than their macroviscosities, while the Dextran10 and PEG8000 microviscosities are comparable to their macroviscosities (Figure S7).

**Figure 2.**
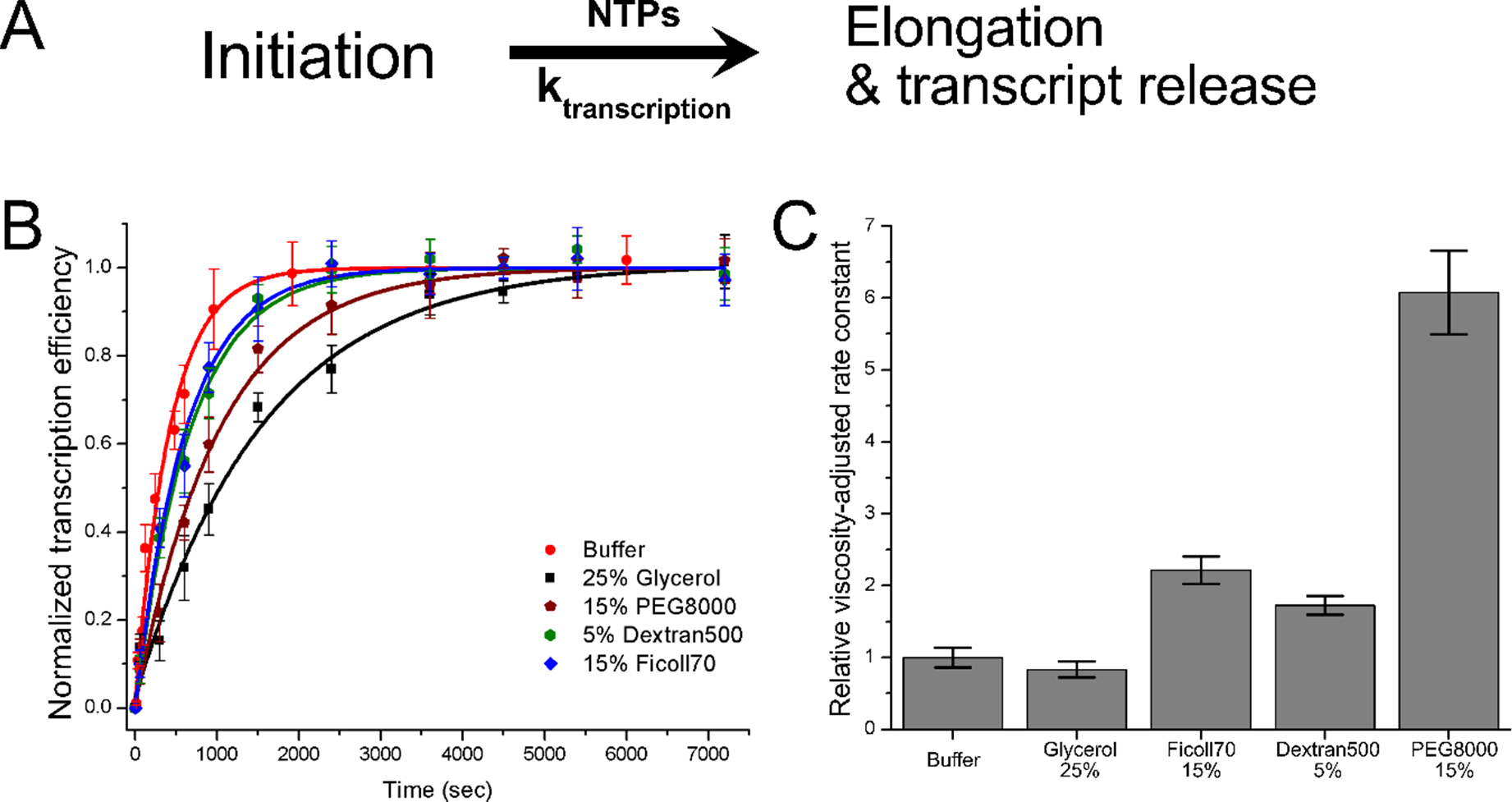
Transcription kinetics at different osmolyte conditions measured by the *in vitro* single-round quenched kinetics transcription assay. (A) Simplified transcription model used for extraction of kinetic constants from the results of *in vitro* quenched kinetics assays. (B) Transcription kinetics in the presence of 25% glycerol (black), 15% PEG 8000 (wine), 5% Dextran500 (olive), and 15% Ficoll70 (blue) compared to transcription kinetics in buffer (red). All transcription efficiencies were normalized to the reaction at 25min without a crowder. (C) The relative viscosity-adjusted rate constants (compared to the rate constant without a crowder). In (B), Error bars in (B) represent the standard deviation of triplicate repeats. Error bars in (C) were calculated based on microviscosities for osmolytes and standard errors obtained from the fitting procedure.

To better understand the effects of osmolytes of various sizes on transcription kinetics, a unidirectional first-order kinetics model (Fig. 2A, See SI for the details) was used to fit and extract kinetic rate constants. To isolate the osmolyte excluded-volume contribution to transcription kinetics, we adjusted extracted rates by the viscosity using the Kramers kinetic theory (which inversely relates rate constants to viscosity)(8, 9, 12, 30–32). This viscosity-adjustment decouples the effect of viscosity from crowding effects due to volume exclusion.

Surprisingly, the viscosity-adjusted rate constants exhibit a significant acceleration by a factor of 2 (Dextran500 & Ficoll70) or 6 (PEG8000) in transcription kinetics for large size osmolytes (Fig. 2C). On the other hand, the viscosity-adjusted kinetics was only marginally affected by 25% glycerol (as compared to reaction in buffer) as expected for glycerol, known as a viscogen that enhances only the viscosity of the medium.

Since all transcription kinetics measurements exhibited first-order (exponential) kinetics (Fig.2B) and same steady-state levels for all crowding conditions (regardless of osmolyte type), it was possible to study the effect of osmolyte sizes and concentration by simpler (and higher throughput) single-time-point transcription assays (transcription reaction quenched at a single constant-time point).

Transcription reactions were initiated by incubation with all NTPs for a constant time of 15 minutes for various osmolyte sizes and concentrations. A 15-minute single time point was chosen since it exhibited the highest sensitivity for changes (as function of osmolytes) in transcription efficiencies (Fig. 2B). Since the reaction conditions were supposed to be for a single round of transcription starting from an RNAP bound to a promoter DNA, the assay examines how osmolytes affect initiation and promoter clearance (*i.e.* excluding the RNAP-promoter binding step).

Our results show that regardless of osmolyte size, the number of transcribed RNAs at a given time (as compared to an identical reaction with no osmolyte) decreases with increase in the osmolyte concentration. However, this reduction is not due to decreased RNAP activity. Rather it is due to the osmolyte modulating the kinetics of the transcription reaction (note that all kinetic curves reach the same steady-state transcript number, Fig. 2B). This result is expected because according to Kramers theory kinetics depend reciprocally on viscosity (8, 9, 12, 30–32).

If the effect of osmolytes was only to increase viscosity, the same transcription efficiency should have been measured for all osmolyte sizes at the same viscosity. We found that transcription efficiencies at a constant viscosity are similar for small PEGs (*i.e.* no size dependence for Ethylene glycol (EG), PEG 200, and PEG 400, similar to glycerol, see bottom 4 curves in Fig. 3B). For larger osmolytes, however, reductions in transcription efficiency (at a constant viscosity) were size dependent (Fig. 3B & 3C). For large PEGs (PEG600~PEG8000), the reduction in transcription efficiency was noticeably smaller (as compared to that of small PEGs) and exhibited an inverse relation with respect to size (top 4 curves in Fig. 3B).

**Figure 3.**
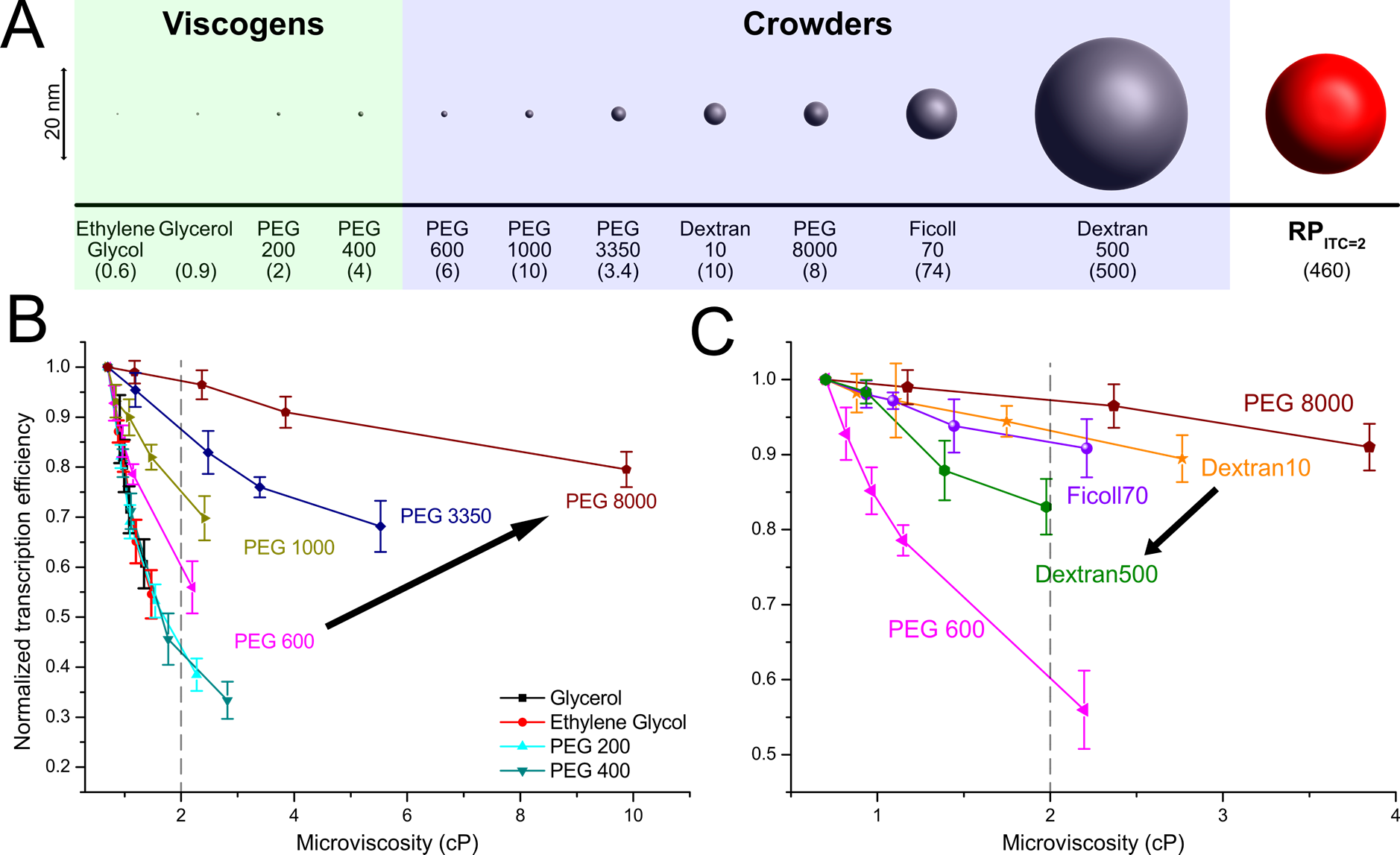
Normalized transcription efficiencies (at a single time point, t_incubation_= 900s) for different osmolytes, as a function of microviscosity. (A) Sizes of osmolytes used in this study. The sizes of osmolytes were obtained from ref.(33–36) and RP_ITC=2_ size was estimated by FCS measurements (see SI for details). The average molecular weights (in kDa) appear in the parentheses. (B, C) Transcription efficiencies measured by single-time-point transcription-quenched kinetics assays for various sizes of PEGs (B) and for Dextran & Ficoll (C). Transcription efficiency values are normalized to the reaction without osmolytes (buffer only). Error bars represent standard deviations of triplicate runs. Lines are guides to the eye. The black arrows in (B) and (C) indicate the direction of increase in the osmolyte size.

To corroborate these results, we performed additional transcription assays with RNA binding dye. This assay showed similar trends distinguishing between small (e.g., Glycerol and PEG 400) and large (e.g., PEG 3500 and 8000) osmolytes (Fig. S5).

Large osmolytes, Dextran and Ficoll, showed a reduction in transcription efficiency that lies between those of PEG600 and PEG8000 (at a given viscosity). For a given microviscosity value, the larger the osmolyte the larger the reduction in transcription efficiency (Fig.3C). Interestingly, this trend is opposite to the trend for increased PEG sizes (Fig. 3B & 3C).

The crowding effect stems from volume exclusion. We therefore plotted in Figure 4 the viscosity-adjusted rate constants (*k*) as function of osmolyte volume occupancy (ϕ_*c*_). To estimate the rate constants while decoupling viscosity effects from the data acquired at a single time point (900 sec), we derived an equation (Eq. S6) based on Kramers kinetic theory and a 1^st^-order kinetics model (Fig 2A). Eq. S4 was used to calculate volume occupancies.

**Figure 4.**
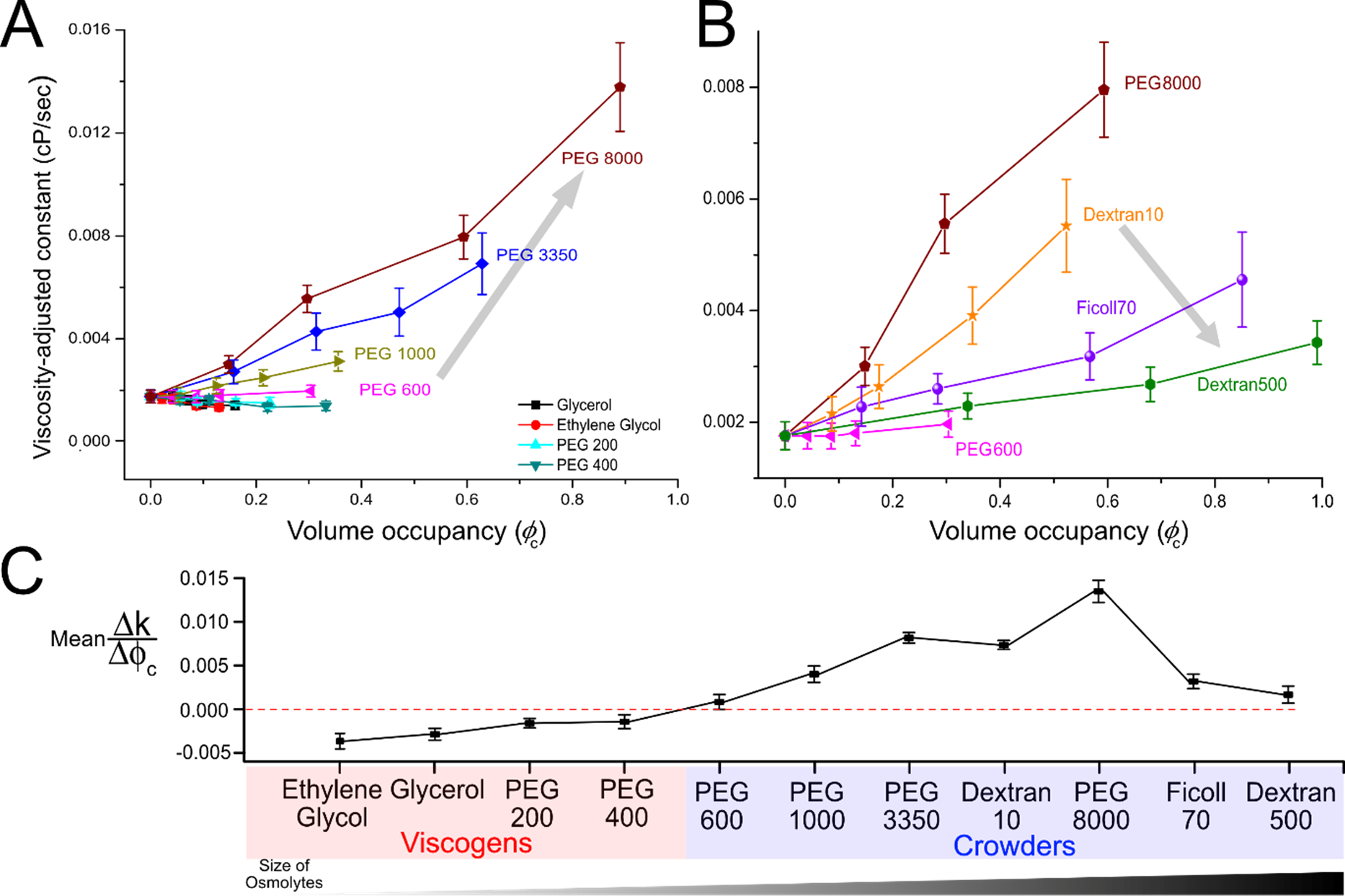
Viscosity-adjusted rate constants as a function of osmolyte volume occupancy. Viscosity-adjusted rate constants were calculated according to Eq.S6 (using results reported in Fig. 2). (A) Small PEGs (< PEG 600) do not affect transcription kinetics (excluding effect of viscosity). Large PEGs (PEG 600 ~ 8000) show strong dependence on size and volume occupancy. (B) Although larger osmolytes (Dextran10& 500, Ficoll70) show dependence on size and volume occupancy, the trend of size dependence is opposite to that for large PEGs. Errors are propagated according to Eq. S6. (C) Linear fits to all curves in (A) and (B) against osmolytes were performed to show qualitatively the trend of transcription kinetics as a function of osmolytes. Lines are guides to the eye. The transparent grey arrows indicate the direction of increase in the sizes of osmolytes.

We found that small PEGs affected transcription rate constants (as a function of volume occupancy) only very marginally (Figs. 4A and 4C), *i.e.* they do not act as crowders, but rather as viscogens (they only increase the viscosity of the medium). In contrast, large osmolytes (large PEGs, Dextran, and Ficoll70) strongly affected transcription rate constants. The rate constants depended on size as well as on volume occupancy of the osmolytes, therefore identifying them as crowders. Interestingly, we found that the dependence of transcription kinetics dependence on large PEGs is opposite in sign to that of larger crowders such as Dextran and Ficoll, although both types showed an overall acceleration of transcription that is proportional to volume occupancy (Figs. 4B and 4C).

Another interesting crowding-related phenomenon was observed for high concentrations of PEG8000 whereby RNAP-Promoter complexes reversibly aggregated (Fig. 5). FCS measurements for RNAP-Promoter complexes with added PEG8000 at concentrations of 7.5% and above exhibited:

(i) Exceptionally long residence times τ_D_. Despite the fact that the macroviscosity of 15% PEG8000 and 30% Dextran10 are comparable, the residence time for 15% PEG8000 was much longer (compare blue and green curves in Fig. 5B);

**Figure 5.**
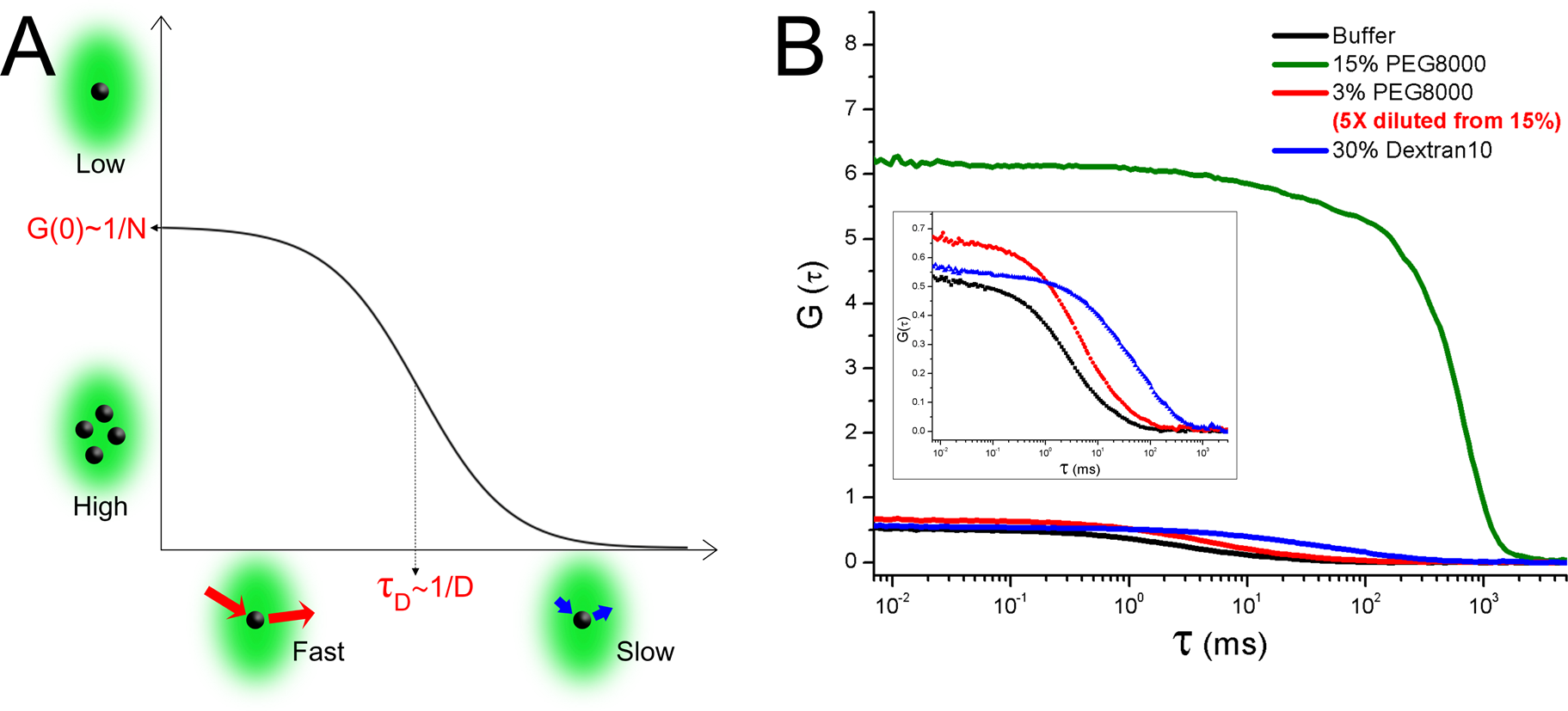
FCS measurements show that RNAP-Promoter complexes could form reversible aggregates in the presence of PEG8000. (A) A schematic of a fluorescence autocorrelation curve for a single species in a solution. The amplitude of the correlation curve extrapolated to zero time delay (*i.e.* G(0)) is inversely proportional to the mean number of fluorescent molecules in the detection volume. The characteristic resident time in the detection volume 1D is inversely proportional to the diffusion coefficient of the molecules. (B) FCS curves for RP_ITC=2_ in buffer (black), 15% PEG8000 (olive), 3% PEG8000 (red) obtained from 5X dilution of 15% PEG8000 whose correlation curve is shown in olive, 30% Dextran10 (blue). Zoomed-in curves for buffer (black), 3% PEG8000 (red), and 30% Dextran10 (blue) are shown in the inset.

(ii) Large correlation amplitude at zero time delay. G(0), the correlation amplitude for 15% PEG8000, is ~ 10 times higher than for all other tested conditions (compare blue, black, and green curves in Fig. 5B). Once the RP_ITC=2_ solution containing 15% PEG8000 was diluted to 3% PEG8000, these features disappeared (compare green and red curves in Fig. 5B). The detailed mechanism for this aggregation is of great interest since previous studies have demonstrated that macromolecular crowding could cause a phase separation that controls enzymatic activities(20, 37, 38). Surprisingly, this aggregation did not affect transcription rates and yields. We plan to explore this regime further in a future study.

## Discussion

We have used an *in vitro* single-round quenched-kinetics transcription assay(26, 27) to investigate the kinetics of *E. coli* RNAP transcription reactions starting from the stable open complex RP_ITC=2_, in the presence of various osmolytes. Our measurements focused on the combined kinetics of abortive initiation, promoter clearance, and elongation. They excluded the kinetics of promoter binding and bubble opening since crowders were added only after bubble formation was established and since very low concentrations of RNAP-DNA complexes were used (preventing re-association of RNAP to DNA, see Methods and Materials for details). We note that the effect of crowding on the RNAP-DNA association reaction is well documented(20, 22).

We found that the activity of RNAP (starting from RP_ITC=2_) is negligibly affected by macromolecular crowding (since kinetic curves for all crowders reached the same steady-state transcription level as in buffer) whereas transcription reaction *kinetics* are strongly affected by macromolecular crowding.

The first effect of crowding is to increase viscosity, which in turn slows the reaction kinetics as predicted by the Kramers kinetic theory(8, 9, 12, 30–32). We found that the microviscosities for large osmolytes such as Ficoll70 and Dextran500 are much lower than their bulk viscosities. In agreement with FCS microviscosity measurements, we found faster apparent kinetics for Ficoll70 or Dextran 500 than for PEG8000 (Fig. 2 and Fig. S7).

The second, and the more interesting observation, is the non-monotonic dependence of reaction kinetics on osmolyte sizes. When viscosity effects are decoupled, small osmolytes such as glycerol and low-molecular-weight PEGs exhibit a negligible effect on transcription kinetics *and thus act solely as viscogens*. Large osmolytes such as Dextran, Ficoll, and high-molecular-weight PEGs, however, exhibit an acceleration in transcription kinetics as their volume fractions increase *and act both as viscogens and as crowders*. Interestingly, while all large osmolytes accelerate transcription kinetics, the larger the PEG the larger the acceleration, while Dextran and Ficoll exhibit an opposite trend: the acceleration for Dextran (500 kDa) is smaller than the acceleration for Ficoll (70 kDa) or Dextran10 (10 kDa).

According to the transcription kinetic model(19, 39–42) (shown in Figure S4), promoter escape is the rate-limiting step of transcription (excluding promoter binding and bubble-opening steps). We hypothesize that the acceleration of the reaction kinetics by crowders is caused by an induced conformational change (to a more compact conformation) in the RNAP-Promoter initial transcribing complexes, resulting in a reduction in the activation barrier for promoter escape.

Scaled-particle theory (SPT) based on the hard sphere model has been commonly and successfully used to explain the effect of macromolecular crowding in terms of volume exclusion (Figs. 6A - 6C)(10, 28). This theory estimates thermodynamic properties of a reaction based on the energy cost for creating a cavity whose size is the same as the one of a newly introduced molecule. According to SPT, the energy gained by compacting a newly introduced molecule is reduced as crowder size increases because larger crowders have larger interstitial cavities. These larger cavities could accommodate expanded molecules more easily. SPT therefore predicts that larger crowders would exclude less volume at a fixed volume occupancy as compared to smaller crowders (Figure 6A - 6C).

**Figure 6.**
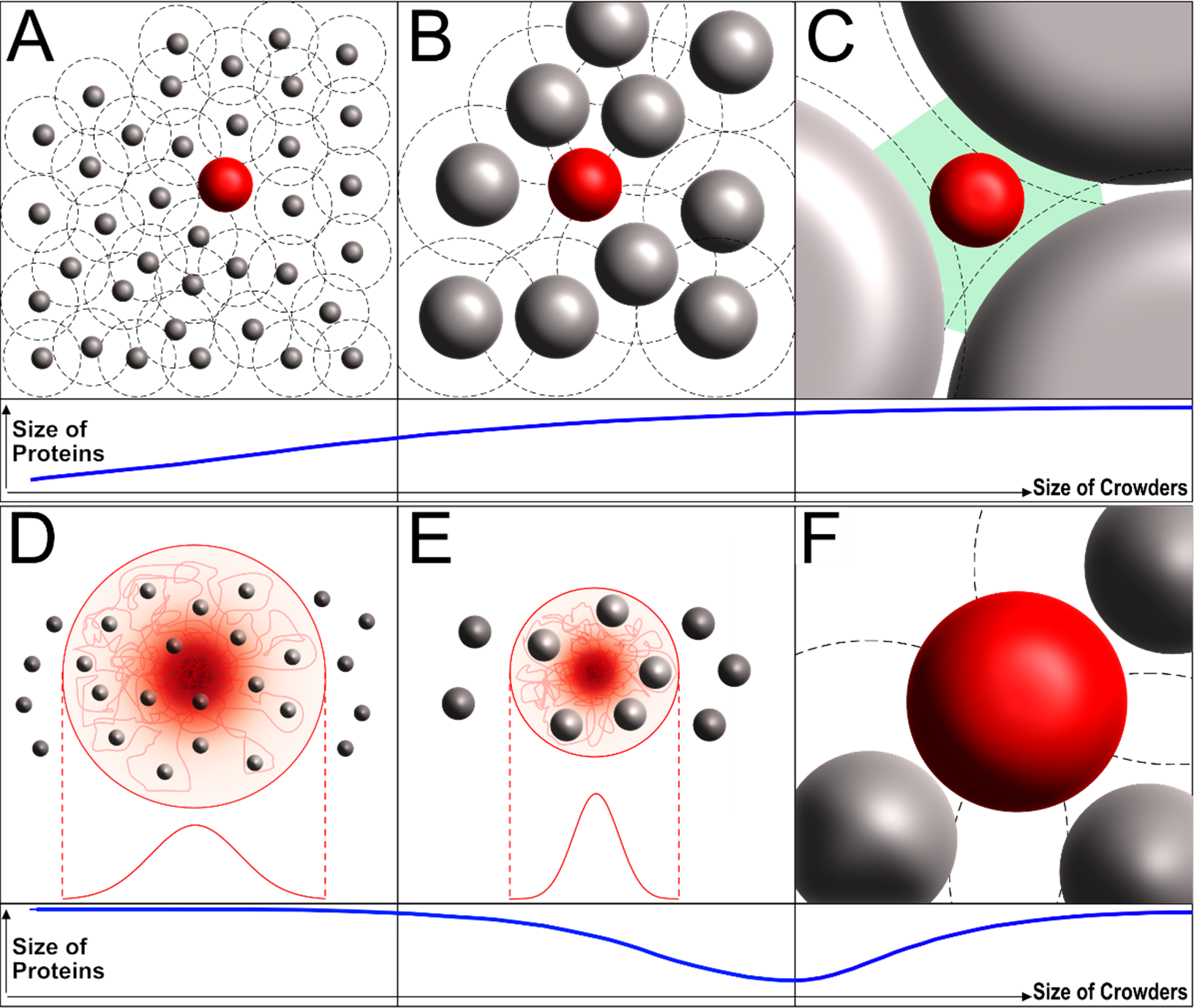
The effect of crowding according to the scaled particle theory (SPT, top) and Gaussian cloud model (bottom). (A – C, top) The Scaled particle theory (SPT)-based hard-sphere model predicts that an unfolded protein chain (red sphere) in the presence of macromolecular crowding becomes less compact as the size of crowder (grey spheres) increases. (A) When the size of a crowder is small compared to the size of the protein, the excluded volume (the volume not accessible to the protein chain, black dashed circle) is much larger than the volume fraction of the crowder. This leads to the compaction of the protein. (B) As the size of the crowder increases, the excluded volume approaches the volume of the crowder, resulting in less compaction (compared to a smaller crowder at the same volume fraction). (C) When the size of the crowder is much larger than the protein, interstitial cavities (green area) can accommodate the unfolded protein chain without compaction so that it will have the same size (or same protein conformation) as in buffer. (D – F, bottom) Gaussian cloud model (GCM) predicts a non-monotonic trend for the size of a protein chain under crowding. (D) A crowder smaller than a given (lower-limit) size would seldom affect the size of the protein (since it permeates the chain, red line). (E) As the crowder size increases (above the limit in (D)), the less volume would be available for the protein chain (red line) since the chain can no longer occupy the same volume with the rigid crowders (grey sphere) in the protein cloud. This induces a compaction of the protein. (F) As the size of the crowder increases further, the GCM eventually turns into the SPT (i.e., the crowder is no longer in the cloud of residues).

With our RNAP conformational change hypothesis, SPT would predict a decrease in transcription rates for larger crowders, since larger crowders will not compact RNAP-Promoter complexes by much (as compared to smaller crowders, at a constant volume occupancy). This is indeed the case for Dextran500 & Ficoll70 (Fig. 4B & 4C). We observed, however, an opposite trend for PEGs, which cannot be explained by SPT. This disagreement could be due to failure of the hard sphere description of polymer crowding, which fails when (i) the shape of the crowder or of the biomolecule deviates significantly from a sphere(10) or when (ii) the crowder *penetrates* the biomolecule (i.e. the biomolecule can no longer be approximated by a hard sphere)(14, 28).

Generalizations of hard-sphere models have been proposed to account for such cases(10, 14, 28). The Gaussian cloud model (GCM), proposed by Minton(28), takes into account the fact that a crowder can permeate a biomolecule. In this model, an unfolded protein is treated as a time-averaged spherically-symmetric cloud of residues whose density distribution is described by a Gaussian function centered on the protein’s center of mass (Figs. 6D and 6E), with the rigid (hard-sphere) crowder allowed to penetrate the cloud of protein residues (Figs. 6D and 6E). If the crowder size is small, the density distribution of protein residues would be only slightly affected (Fig. 6D). As the size of the crowder increases, it becomes harder for the unfolded chain to avoid crowders and it will therefore compact itself (Fig. 6E). As the size of the crowder increases further, however, the size of the crowder is too big to be in the protein cloud(28), and the GCM converges to SPT (Fig. 6F). Note that this *non*-monotonic effect (on protein size) of increasing crowder size is not seen in the SPT progression depicted in panels 6A, B, and C.

Applied to the transcription reaction, the GCM would predict that small crowders will have little effect on transcription kinetics. As the crowder size increases (above a lower limit), the transcription kinetics would accelerate. The kinetics, however, will stop accelerating and even start to decelerate once the crowder’s size transitions to the SPT regime, This predicted non-monotonic dependence (Fig. 6, D-F) is consistent with our observations (Fig. 4C). Because GCM deviates from SPT only when the size of the crowder is small, it is worth noting that PEG8000 is much smaller than the RNAP-DNA complex (Fig. 3A, Table S1).

The polymeric nature of a protein suggests that it could be partially or fully infiltrated by a crowder, unlike an interaction with a hard sphere. Although a previous report demonstrated that the T7 DNA polymerase complex became more compacted under macromolecular crowding conditions(37), it is still possible that the size of the entire RNAP-Promoter complex will be only nominally affected by crowders since the majority of the complex is well-structured. However, the complex does contain a few partially unstructured domains, or hinged structured domains that are crucial for the enzyme’s functionality. For example, the trigger loop is partially disordered and its folding propensity determines the rate of polymerization throughout all transcription stages (after promoter open-complex formation)(43, 44). σR3.2 forms an unstructured σ finger loop and its displacement is known to be a common rate-determining step for promoter escape(25, 26, 45). Also, recent studies (25, 26) have suggested that RNAP could be paused in late initiation stages and this pause could be controlled by σR3.2. We therefore argue that macromolecular crowders could possibly compact partially disordered / hinged domains of RNAP that in turn affect transcription kinetic rates. Future studies should examine whether the entire RNAP-DNA complex is affected by crowders, or identify which of the RNAP domains is most affected.

Finally, we emphasize that since the cellular milieu contains various size of crowders such as proteins, glycans, and oligonucleotides, our observation of non-monotonic dependence of crowders’ size on the transcription reaction would provide valuable insight into elucidating how the crowded cell environment plays a role in the transcription process (specifically in regulatory pathways) in the cell.

## Methods and Materials

### 1. Design of the template DNA and probe for *in vitro* single-round quenched kinetics transcription assays.

The DNA template used for the transcription reaction was designed to produce a transcript containing a sequence complementary to the probe sequence (Fig. S1). Thus, an RNA produced by the transcription reaction would be efficiently hybridized with the ssDNA probe. The transcription detection probe is a doubly labeled ssDNA with a donor-acceptor FRET pair - a donor 5-carboxytetramethylrhodamine (TMR) at its 5’ end and an acceptor (Alexa Fluor 647) at its 3’ end. In order for efficient hybridization with transcripts, we designed the ssDNA probe with 20 consecutive deoxythymidines (20 dT) to be unstructured (i.e. with no stable secondary structure) in solution(26, 27). Since the persistence length of unstructured ssDNA is short (~1.5 nm in 2 M NaCl, ~3 nm in 25 mM NaCl)(46), the two dyes are close to each other, yielding a single high FRET population with a peak FRET efficiency of E ~ 0.8. However, when hybridized to the run-off mRNA transcripts, the probe-mRNA assumes a rigid dsDNA-RNA conformation (persistence length for dsDNA ~50nm(47), for dsRNA ~64nm(48)), and the distance between the two dyes is significantly increased, leading to the appearance of a second FRET sub-population with a peak FRET efficiency of ~0.3.

### 2. Preparation of a stable RNAP-Promoter open complex (RP_ITC=2_).

Double-stranded DNA (dsDNA) template, *lac*CONS-GTG-20dA (Fig. S1), was prepared by hybridizing single-stranded DNAs (ssDNA, Integrated DNA Technologies, Coralville, IA, USA). The strands were hybridized in hybridization buffer (40 mM Tris-HCl, pH 8, 150 mM Magnesium Chloride (MgCl_2_)) with a thermocycler (with temperature increased to 95 ºC and then slowly decreased to 21 ºC).

RNAP open complex (RP_O_) was prepared by incubating 3 μL *E. coli* RNAP holoenzyme (NEB, Ipswich, MA, USA, M0551S; 1.6 μM), 10 μL 2X transcription buffer (80 mM HEPES KOH, 100 mM KCl, 20 mM MgCl_2_, 2 mM dithiotreitol (DTT), 2 mM 2-mercaptoethylamine-HCl (MEA), 200 μg/mL Bovine Serum Albumin (BSA), pH 7), 1 μL of 1 μM *lac*CONS-GTG-20A promoter and 6 μL of water at 37 ºC for 30 minutes. To remove nonspecifically-bound RNAP-DNA complexes and free RNAPs, 1 μL of 100 mg/mL Heparin-Sepharose CL-6B beads (GE Healthcare, Little Chalfont, Buckinghamshire, UK) was added to RP_O_ solution together with 10 μL of pre-warmed 1X transcription buffer followed by 1 minute incubation at 37 ºC and centrifugation for 45 seconds at 6000 rpm to pellet Heparin-Sepharose. 20 μL of the supernatant containing RP_O_ was transferred to a new tube, and 1.5 μL of 10 mM the dinucleotide primer Adenylyl(3'-5') adenosine (A_p_A, Ribomed, Carlsbad, CA, USA) was added to the RPo solution, which was then incubated at 37 ºC for 20 minutes to form RP_ITC=2_, stabilizing the open complex. The RP_ITC=2_ solution was used as a stock for all transcription reactions. 2 μL of RNase inhibitor (NEB, Ipswich, MA, USA, M0314S) was added to the solution to protect transcribed RNA molecules from possible degradation.

### 3. *In vitro* single-round quenched kinetics transcription assays

Prepared stock solution of RP_ITC=2_ was diluted in transcription buffer containing a given concentration of the osmolytes. Transcription reactions were initiated by 100 μM of high purity NTPs (GE Healthcare, Little Chalfont, Buckinghamshire, UK) at 37 ºC. The transcription reaction was stopped at a given time ***t*** by addition of 1.5X volumes of quencher/probe solution containing 1.25 M Guanidium Chloride (GdmCl) and 250 pM ssDNA probe (final concentration of GdmCl and ssDNA probe were 500 mM and 100 pM, respectively). Then, the quenched-probed solution was incubated for additional 1 hour at room temperature in order to establish complete hybridization between transcribed RNA molecules and ssDNA probes. The numbers of hybridized probes (to transcripts) and non-hybridized probes are accurately determined using alternating-laser excitation (ALEX)-based fluorescence-aided molecule sorting (ALEX-FAMS)(29, 49).

Experiments were performed with a low concentration of FRET probe (~100 pM) (within the optimal concentration range for ALEX-FAMS). This is a much lower concentration than the RNAP-Promoter binding affinity (K_d_ = 10~100 nM at physiological ionic strength(50)), thereby ensuring that only a single-round run-off transcription reaction occurs on any given template (Re-association of RNAP to a template dsDNA is unlikely after it dissociates from the template). This, in turn, allows us to quantify transcription efficiency under various conditions or at different times after initiation of the reaction by simply counting the number of transcripts. We could therefore study how osmolytes affect the efficiency of transcription for varying concentrations and the effect of sizes and types of different osmolytes.

Although the same amounts of reagents were used to generate the RNAP-Promoter open bubble, the efficiency of formation could vary for many reasons. In this context, the concentration of the RNAP-DNA complex was calibrated in order for transcription efficiency at equilibrium (t=∞) not to exceed the dynamic range of the transcription assay (The concentration of transcripts cannot be more than that of the probe (~100pM) at equilibrium).

For all *in vitro* quenched kinetics transcription assays, data were acquired for 15-20 minutes using the ALEX-FAMS setup, as described in Kapanidis et al.(29) with two single-photon Avalanche photodiodes (SPADs, Perkin Elmer Inc., Waltham, MA, USA) and 532 and 638 nm CW lasers (Coherent Inc., Santa Clara, CA) operating at powers of 170 and 80 μW, respectively. All FRET data analyses in this work have been done using a Python-based open-source burst analysis toolkit for confocal single-molecule FRET, FRETBursts(51) and a Python package for non-linear least-squares minimization and curve-fitting, lmfit(52).

### 4. mRNA detection by RNA binding dye assay

For 50 μL reactions, prepared stock solutions of stable RP_ITC2_ were diluted to the final concentration of 1 nM (based on the concentration of promoter DNA) in transcription buffer containing different types and concentrations of crowders. Transcription reactions were initiated by 100 μM of NTPs and incubated at 37 ºC for 15 minutes. DNase I was added with DNase buffer (NEB, Ipswich, MA, USA, M0303S for DNase I and M0303S for DNase reaction buffer) to each sample, resulting in a total volume of 100 μL, to digest the DNA template and stop the transcription reaction at 37 ºC for 1 hour. Pre-prepared 100 μL of Quant-iT™ OliGreen^®^ solution (Thermo Fisher Scientific Inc, Canoga Park, CA) was then added to each sample to measure the amount of mRNA produced by transcription reactions. Fluorescent signals were obtained with a Tecan Infinte M1000 Plate Reader (excitation wavelength 480 nm. emission wavelength 520 nm)

### 5. Microviscosity measurements using Fluorescence Correlation Spectroscopy (FCS)

We performed FCS measurements to measure microviscosity for RNAP-Promoter complexes under given crowding conditions, *lac*CONS-GTG-20dA promoter labeled with Atto550 at −5 register on the non-template strand and Atto647N at −8 register (in reference to the transcription start site) on the template strand(26, 53) was used for preparation of dual-labeled stable RNAP-Promoter complexes (RP_ITC=2_) as described in #2 Above. The dual-labeled RP_ITC=2_ solution was analyzed by ALEX-FAMS before the FCS measurements to ensure that the final solution contains mostly (≥ 90%) RP_ITC=2_ (and not free promoter DNA, see Fig. S6).

All FCS measurements were performed at 25 °C using a home-built confocal microscope based on Olympus IX71 with CW laser at 638nm (Melles Griot Inc., Carlsbad, CA). The emitted photon stream from the labeled complexes was equally split into two SPADs (Perkin Elmer Inc., Waltham, MA, USA). Detected fluorescence signals were sent to ALV-6000 MultiCorr digital real time correlator then cross-correlated to eliminate detector after-pulsing. Each sample was measured three times.

## Supporting Information

Supporting information is available on the website. The information provides further details on analysis and validation of transcription assays, microviscosity estimation from FCS measurements, volume occupancy estimation, kinetic model, derivation of Eq. S6, additional figures showing the sequences of the *lac*CONS-GTG-20dA and ssDNA probe used in this study, the results of the RNA binding assay and microviscosity measurements. Hydrodynamic radii of the crowders used in this study appear in Table S1.

## AUTHOR INFORMATION

### Notes

The authors declare no competing financial interests.

## ACKNOWLEDGMENT

The authors thank Dr. Xavier Michalet and Dr. Antonino Ingargiola for fruitful scientific discussions and consultations. The authors also thank Maya Lerner for helping on preparation of illustrations. This work was supported by the NIH (GM069709 to SW) and NSF (MCB-1244175 to SW, and CHE-1051507 to WMG and CMK).

